# Predicting the effects of multiple global change drivers on microbial communities remains challenging

**DOI:** 10.1101/2021.12.13.472339

**Authors:** Marcel Suleiman, Uriah Daugaard, Yves Choffat, Xue Zheng, Owen L Petchey

## Abstract

Microbial communities in many ecosystems are facing a broad range of global change drivers, such as nutrient enrichment, chemical pollution, and temperature change. These drivers can cause changes in the abundance of taxa, the composition of communities, and the properties of ecosystems. While the influence of single drivers is already described in numerous studies, the effect and predictability of multiple drivers changing simultaneously is still poorly understood. In this study, we used 240 highly replicable oxic/anoxic aquatic lab microcosms and four drivers (fertilizer, glyphosate, metal pollution, antibiotics) in all possible combinations at three different temperatures (20 °C, 24 °C, and 28 °C) to shed light into consequences of multiple drivers on different levels of organization, ranging from species abundance to community and ecosystem parameters. We found (i) that at all levels of ecological organisation, combinations of drivers can change the biological consequence and direction of effect compared to single drivers (ii), that effects of combinations are further modified by temperature, (iii) that a larger number of drivers occurring simultaneously is often quite closely related to their effect size, and (iv) that there is little evidence that any of these effects are associated with the level of ecological organisation of the state variable. These findings suggest that, at least in this experimental ecosystem approximating a stratified aquatic ecosystem, there may be relatively little scope for predicting the effects of combinations of drivers from the effects of individual drivers, or by accounting for the level of ecological organisation in question, though there may be some scope for prediction based on the number of drivers that are occurring simultaneous. A priority, though also a considerable challenge, is to extend such research to consider continuous variation in the magnitude of multiple drivers acting together.

## Introduction

Microbial communities are key components of many ecosystems, and are often exposed to many anthropogenic global change drivers (Christensen et al. 2006; Jackson et al. 2016). Exposure to fertilizer (da Costa et al. 2021) and metal pollution (Xu et al. 2018a), pesticides like glyphosate (Solomon and Thompson 2003; Relyea 2009), antibiotics (Xu et al. 2018b) and temperature increase (Wu et al. 2011) force populations and whole ecosystems to develop in different ways compared to less or unaffected ones (Tylianakis et al. 2008). Studies show that even slight changes in environmental conditions can lead to large taxonomic and functional microbial change, and even to regime shifts, with potential long-term consequences for biogeochemical cycles and ecological function of the affected habitat (Gruber 2011).

While many studies have focused on one global change driver, recent studies (Christensen et al. 2006; Rillig et al. 2019; Suleiman et al. 2021a) indicated that the combined effect of multiple drivers applied simultaneously is often not equal to the sum of the effects of each individual driver. Rather, they demonstrate that interactive effects can occur. These can be synergistic (combined effect greater than the sum of the effect of individual drivers) or antagonistic (combined effect less than the sum of the effect of individual drivers). Furthermore, not just the type of driver, but also the number of factors combined plays a crucial role (Rillig et al. 2019). Nevertheless, the number of studies testing three or more drivers remains very low (Rillig et al. 2019), which highlights the need for further investigations of combined effects. Combining this with the importance of understanding drivers of change in microbial communities is a critical aspect of the emerging research field of climate/global change microbiology (Hutchins et al. 2019).

The limited number of studies with three or more drivers may be in part due to the logistical demands of the required experiments and the complexity of interpreting their results. For example, to assess all interactions a fully-factorial experimental design is needed which quickly leads to large experiments. Regarding interpretation, it can be difficult to understand the meaning of interactions among more than three drivers, such that even if one would conduct a fully factorial experiment, traditional methods for interpreting interactions (such as interaction plots) may be insufficient. Some studies have instead focused on the effect of variation in the number of drivers, and have applied only a subsample of all possible combinations (e.g. Brennan and Collins 2015; Rillig et al. 2019).

Another gap in understanding is how drivers and combinations of drivers act across levels of ecological organization from individuals to ecosystems (Galic et al. 2018). For example, in a case study of a model of amphipod feeding behavior Galic et al (2018) showed that responses to multiple drivers at the individual level were not consistent with those at higher levels of organization. In their case-study, the nature of this inconsistency would lead to underestimation of effects at population and ecosystem levels if effects at the individual level were assumed to hold across levels of organisation.

These gaps in knowledge are problematic. One reason is that interactions can be source of ecological surprises, i.e., if we assume additivity we can be surprised if there are interactions (Christensen et al 2006). Also, unless there are some generalities about interactions among drivers, we will never be able to predict the effect of a new (previously unstudied) combination of drivers, so interactions will also then be a surprise (Christensen et al 2006). Hence, we and others (e.g., Simmons et al 2021, Rillig et al 2018) are interested in discovering if there are any general patterns that will allow us to predict the effects of combinations of environmental changes. This report is about our search for signals of general patterns in the effects of combinations of environmental drivers. For example, whether the relationship between a biological variable and the number of drivers is dependent on the level of ecological organization of the biological variable and whether the strength and nature of the interaction effects varies with level of ecological organization.

Microbial communities in aquatic ecosystems are complex (Chistensen et al. 2006; Davis et al. 2010; Faust and Raes 2012), consisting of feedback among numerous abiotic and biotic interactions (Singh et al. 2009), among and within functional microbial groups (Bush et al. 2017; Richardson et al. 2018). Recent work indicates that these ecosystems are sensitive to environmental change (Christensen et al. 2006; Shade et al. 2011, 2012; Suleiman et al. 2021b, a), identifying them as appropriate systems for the study of the influences of global changes.

In this work, we applied four different global change drivers (fertilizer, glyphosate, metal pollution, and antibiotics) together with increasing temperatures (20 °C, 24 °C, and 28 °C) in all combinations possible, on a recently developed and highly replicable stratified aquatic microbial lab system. Our experiment includes the analysis of several abiotic (oxygen, total nitrogen, total organic carbon, pH) and biotic variables (Shannon index, microbial community composition, genera abundances), since recent studies have shown that various levels of ecosystems can be affected (Weithoff et al. 2000; Shade et al. 2011, 2012; Suleiman et al. 2021b) and in order to examine if effects show predictable variation across levels of organisation. We hypothesize that (i) combinations of drivers will have non-additive effects on system properties, (ii) the non-additive effects will be different depending on the temperature, and (iii) increasing the number of drivers applied will cause a systematic change in system properties.

## Material and methods

### Experimental ecosystems

Incubation of stratified microbial communities was performed in microcosms consisting of standard glass test tubes (4 mL volume). These test tubes were closed with plastic lids, with a small hole (0.5 mm) that allowed gas exchange between the headspace and the atmosphere. Each microcosm consisted of sediment and water samples taken in May from a small pond in Zurich, Switzerland (47°23’51.2”N 8°32’33.3”E, temperature of 19 °C, pH of 7) at a depth of 25 cm. Sediment was homogenized (30 min mixing) and subsequently supplemented with sterile 0.25 % crystalline cellulose, 0.25 % methyl-cellulose, 0.5 % CaSO_4_, 0.1% CaCO_3_, and 0.005 % NH_4_H_2_PO_4_. Glass tubes were filled with a 3 mm layer of supplemented sediment and covered with 3.4 mL pond water (with 0.005 % NH_4_H_2_PO_4_), resulting in 500 µL headspace. Microecosystems were incubated at room temperature for 2 hours to let sediment settle, before the different treatments were applied. The experiment involved incubation for 23 days at either 20 °C, 24 °C or 28 °C.

### Experimental design

The four drivers, namely NH_4_H_2_PO_4_-fertilizer (F), Glyphosate (G), metal pollution (M) and antibiotics (A), each had two levels. Furthermore, control microcosms (C) were used without application of any drivers and the experiment was carried out at 20 °C, 24 °C and 28 °C under a dark-light cycle of 8:16 h. The temperature treatment and four driver treatments were applied in a fully factorial design. Five replicates (Burgess et al. 2022) per treatment combination resulted in 240 microcosms, according to the following design:

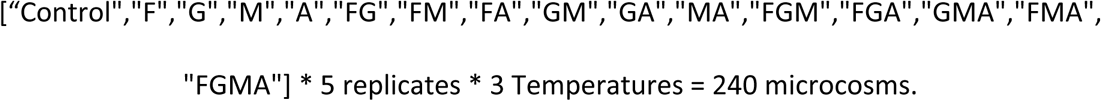

For the driver treatments, the following chemical compounds were added to the microcosms two hours after the assembly of the microcosms:

- NH_4_H_2_PO_4_ in final concentration of 0.1 g/L as fertilizer (F), as used in a previous study (Suleiman et al., 2021)
- Glyphosate in final concentration of 0.001 g/L (G) as used in a recent study (da Costa et al. 2021)
- Metal pollution: DSMZ Trace element solution SL-10 final concentration 10x higher than recommended (https://www.dsmz.de/microorganisms/medium/pdf/DSMZ_Medium320.pdf). The following chemicals with final concentration in the microcosms were 0.015 g/L FeCl_2_ x 4 H _2_ O,. 0.7 mg/L ZnCl_2_, 1 mg/L MnCl_2_ x 4 H_2_O, 0.06 mg/L H_3_BO_3_, 1.9 mg/L CoCl_2_ x 6 H_2_O, 0.02 mg/L CuCl_2_ x 2 H_2_O, 0.24 mg/L NiCl_2_ x 6 H_2_O, 0.36 mg/L Na_2_MoO_4_ x 2 H_2_O (M)
- Mixture of penicillin and ampicillin (0,05 mg/100 mL each) in final concentration of 25 ng/L for antibiotic treatment (A), since comparable maximum amounts were detected in natural aquatic habitats (Xu et al. 2018b)

Drivers were applied to the microcosms only on day 0; there were no further press or pulse perturbations. After incubation for 23 days, the whole microcosm was centrifuged for 5 min at 10’000 rpm (sediment and water column together). The pellet was used for DNA extraction the supernatant was used for performing TOC analysis.

### Measurement of abiotic factors

Oxygen concentration was measured 1.5 cm below the water/air interphase using an oxygen dipping probe (PreSens, Germany) at the last day of incubation. pH was measured in the supernatant of the centrifuged microcosm. Total nitrogen (TN) and total organic carbon (TOC) were measured in the supernatant using a TOC analyser.

### Microbial community composition

DNA was extracted using ZymoBIOMICS DNA Miniprep Kit (ZymoResearch), following the manufacturer’s instructions. Full-length 16S rRNA gene amplification was performed using the primer pair 27F forward primer (5’-AGRGTTYGATYMTGGCTCAG-3’) and 1592R reverse primer (5’-RGYTACCTTGTTACGACTT-3’), as reported previously (Suleiman et al., 2021).

PCR products were visualized on an 1 % (w/v) agarose gel and were pooled for sequencing in equal concentrations. PCR product pools were purified using AMPure®PB beads (PacBio). Sequencing was performed at the Functional Genomic Center Zürich, Switzerland, and performed using SMRT Technology (PacBio) as reported previously (Suleiman et al. 2021a) (on three SMRT chips). Sequencing data quality were checked using the PacBio SMRT Link software.

Sequencing data were filtered based on primer sequences, length (1300-1600 bp), quality, error rates and chimeras using the R-package *dada2* (Callahan et al. 2016). The final sequence table was aligned using SILVA ribosomal RNA database (Quast et al. 2012), using version 138 (non-redundant dataset 99). A phyloseq object was created using the *phyloseq* R-package (McMurdie and Holmes 2013), consisting of amplicon sequence variant (ASV) table, taxonomy table and sample data. For further analysis, the R-packages *phyloseq* (McMurdie and Holmes 2013) and *vegan* (Oksanen et al. 2019) were used. Overview on number of reads per sample after the dada2 pipeline is shown in supplementary table 1.

To compare the effects of the multiple drivers, we analysed various response variables, namely community composition based on non-metric multidimensional scaling NMDS, Shannon index, oxygen, total nitrogen TN, total organic carbon TOC, and abundances of specific genera and functional groups.

Microbial community composition was quantified using NMDS based on Bray-Curtis scores with the *metaMDS* function of the vegan R-package (Oksanen et al. 2019), with three dimensions used (k=3, try=100). We chose to use NMDS to better represent the non-linearities that often occur in ecological patterns that lead some other types of ordination (e.g. principal component analysis) to be less desirable. All samples were analysed at once in the NMDS analysis (Stress = 0.11; stress less than 0.2 is generally considered acceptable, e.g. (Dexter et al. 2018). Shannon index was calculated using the *phyloseq* package (McMurdie and Holmes 2013). We included the analysis of the Shannon index since the system is highly dynamic and develops from a small microbe density to a high microbe density. Please note that we can not differentiate between live and dead as well as active and non-active cells. Functional groups were constructed by filtering for cyanobacterial, sulfur-related and typical lake-associated species, which resulted in a reduced dataset consisting of cyanobacterial orders *Cyanobacteriales, Synechococcales, Leptolyngbyales* and *Limnotrichales*, phototrophic sulfur-bacterial orders of *Chromatiales* and *Chlorobiales*, as well as sulfate-reducing orders of *Desulfobulbales* and *Desulfovibrionales*. Furthermore, we also implemented the order Campylobacteriales, since some members (not all) like *Sulfuricurvum* and *Thiovolum* are also associated with the sulfur cycle. The community of these members were used as a subset for NMDS analysis in this study (stress = 0.10). Detailed scripts and a phyloseq with all sequences are available in the github repository https://github.com/Marcel2907/multiple-stressor and on zenodo (DOI 10.5281/zenodo.5948600).

### Data analysis

For every response variable (see above) we carried out two linear models, one to investigate the relation between the response variable and the various drivers and their combinations and one to investigate how the response depended on the number of drivers applied. To ensure that modelling assumptions were met, we carried out the following response variable transformations, which are listed in detail in the deposited R script. For total nitrogen we took the square root, for total carbon we took the squared inverse, for *Tychonema* we used a logit transformation and for *Sulfuricurvum* we took the cubic root. For the variables not listed here no data transformations were necessary. Further, for total nitrogen concentration we excluded 13 data points due to measurement problems (Supplementary Figure 1). For the analyses of the effect of the number of applied drivers, all response variables were further standardized to render them more comparable. Moreover, in the cases of pH and total nitrogen we included the variable fertilizer as a covariate, as it strongly influenced these response variables. In all models we included all possible covariate interactions.

## Results

### Microbial community and genera abundances

In total, 20’319 unique ASVs were identified after running the dada2 pipeline. After filtering to retain only ASVs with relative abundance of at least 0.1 % in at least one sample, the total number of ASVs decreased to 5’830 taxa. The control microcosms consisted of mainly cyanobacteria (e.g., *Phormidaceae*) and phototrophic sulfur bacteria (specially *Chromatiaceae*) (Fig. 1 a&b, Supplementary Fig. 2). Depending on the driver combination and temperature, the microbial communities and genera abundances showed strong differences compared to the controls. Microcosms with glyphosate and metal pollution, each as a single driver or in combination, were still dominated by members of *Phormidiaceae* and *Chromatiaceae*, but in all remaining treatments the abundance of members of these families was lower (Supplementary Fig. 2).

**Fig. 1.**
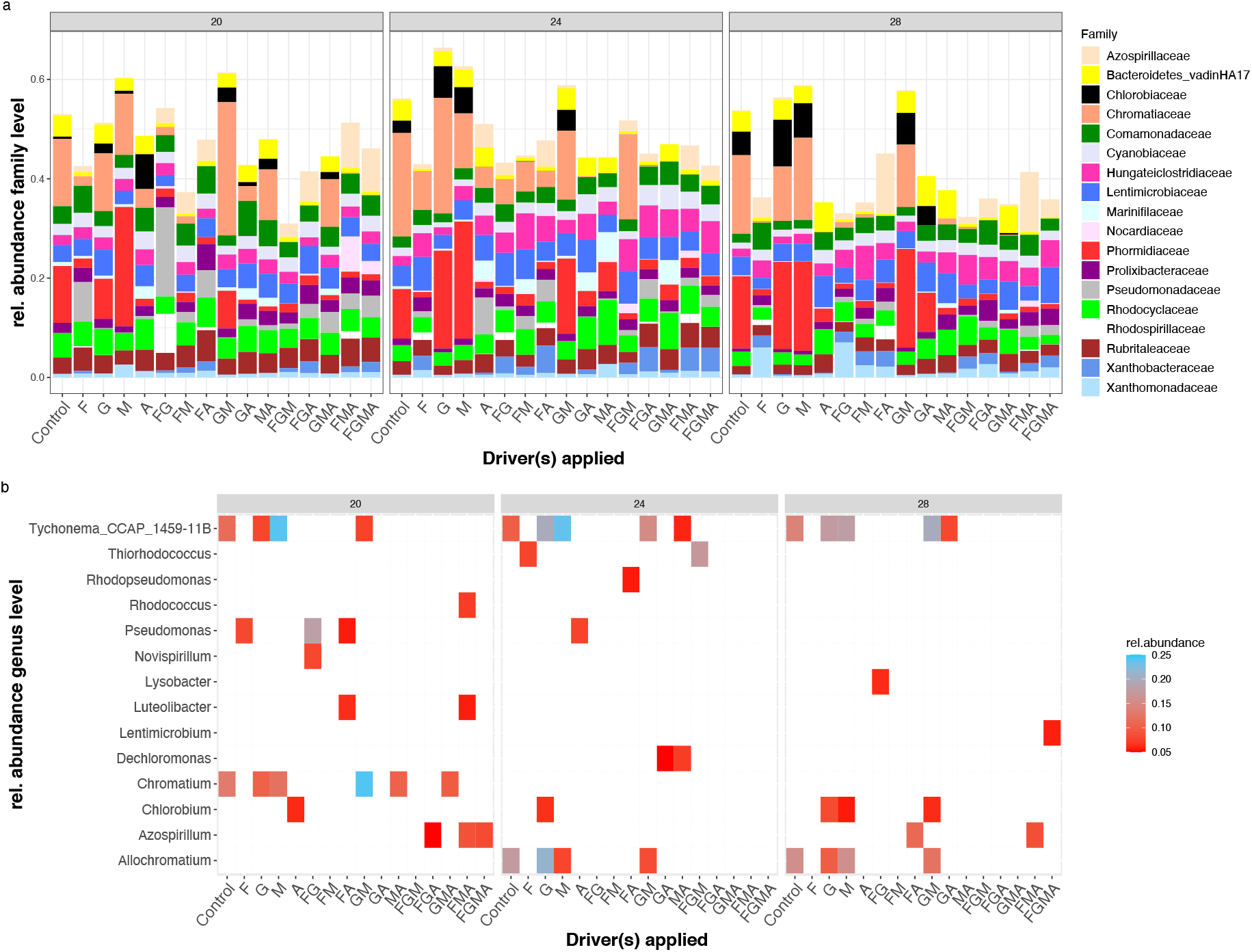
Effects of drivers and driver combinations on microbial community composition and genera abundance. **(a)** Microbial community composition (mean per 5 replicates) at family level. Taxa with rel. abundance > 5 % shown. **(b)** Relative abundance of specific genera with rel. abundance > 5 % for each driver combination at each of the three temperatures. White color represents a relative abundance < 5 %. F=Fertilizer, G= Glyphosate, M=Metall pollution, A=Antibiotics.

Also on genus level, we observed shifts in microbial composition due to driver combination and temperature (Fig. 1 b, Supplementary Fig. 3). The cyanobacterial genus *Tychonema* was highly abundant in the metal pollution only treatment, and with increasing temperature also in glyphosate+metal pollution treatments (20 % at 28 °C, 7 % at 20 °C). *Chromatium*, in contrast, appeared in several treatments and reached highest relative abundance at 20 °C when confronted with G:M (20 %), but vanished with increasing temperature in all treatments (< 5%). Statistical analysis of the dependence of relative abundance on driver combinations and temperature were performed on genus level for *Tychonema* and *Sulfuricurvum*, as representative species for high-abundance and low-abundance taxa (Fig. 3). Statistical analysis confirmed significant changes of relative species abundance caused by individual drivers, due to combination of drivers, and due to temperature.

### Effects of combination and number of drivers on Shannon index

Shannon index of the microcosms changed depending on the combination of drivers (Fig. 2a+b), resulting in values between 3 and 7, with the majority of samples between 5 and 6, indicating diverse microbial communities within the microcosms. Several significant driver treatments and combinations effects occurred, namely temperature increase (F-statistic = 6.8, p-value = 0.001), fertilizer (F-statistic = 16.0, p-value < 0.001), antibiotics (F-statistic = 26.3, p-value < 0.001), as well as combination of fertilizer+antibiotics (F-statistic = 17.9, p-value < 0.001) and fertilizer+antibiotics+glyphosate (F-statistic = 4.8, p-value = 0.030) (Fig. 2a). All significant driver combinations had a positive effect on Shannon index, except the temperature increase to 28 °C and the fertilizer+antibiotic combination treatment (Fig. 3, Fig. 6). Interestingly, the negative effect of fertilizer+antibiotic was reversed to a positive effect again by adding glyphosate, while the treatment of glyphosate alone had no impact (Fig. 2a, Fig. 3). Number of drivers did not affect Shannon index, while temperature did (Fig. 2b, Fig. 4).

**Fig. 2.**
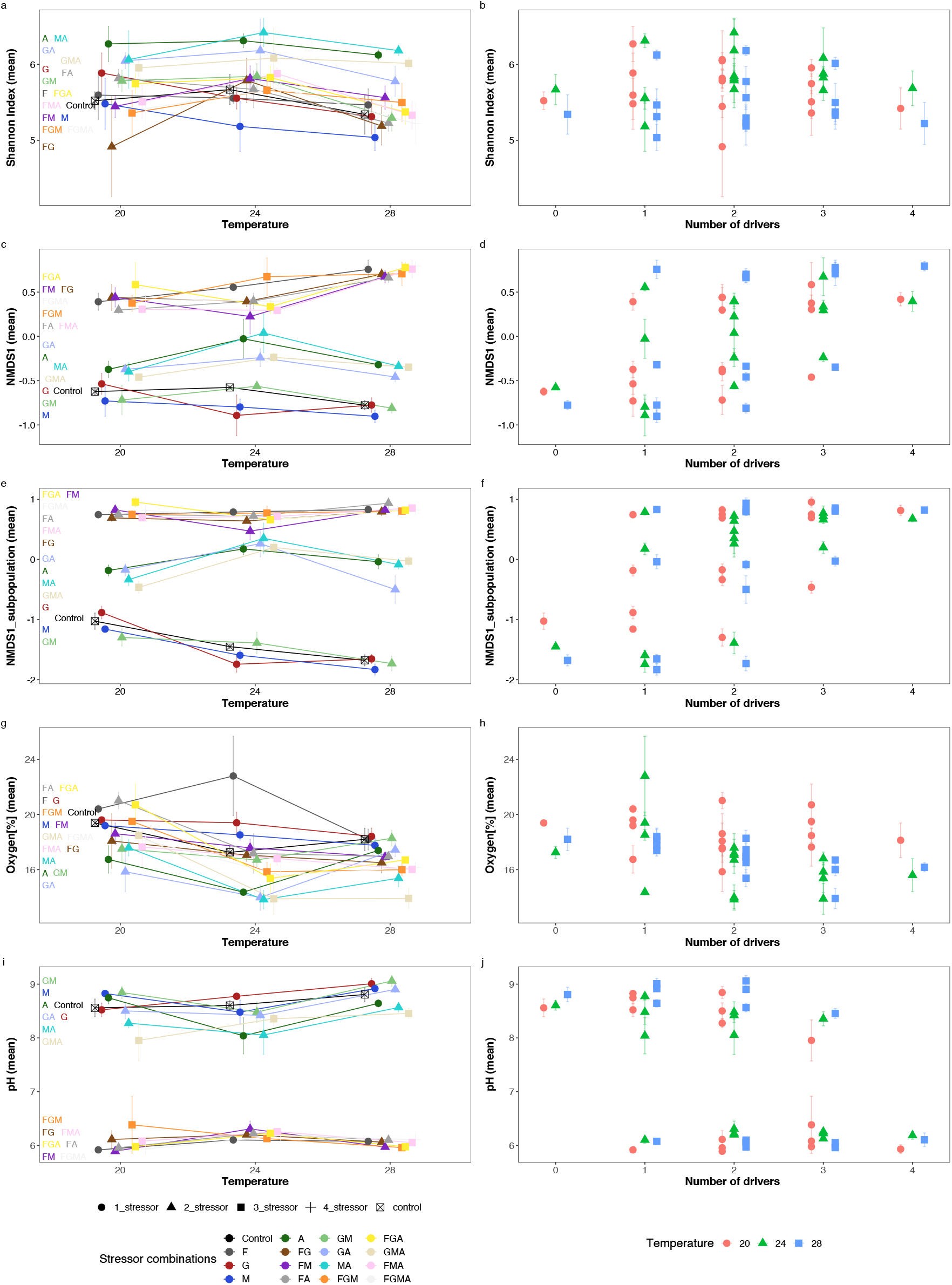
Effects of driver combination, number of drivers, and temperature on microbial community and ecosystem variables. Each row of the figure contains two graphs, both of which concern the same response variable, e.g., first row (a) and (b) concern Shannon index. The left column of figures shows the effects of temperature and driver combination. The right column shows the effects of temperature and number of drivers. Symbols are the mean of the five replicates, and errors bars show the standard error of the mean.

**Fig. 3.**
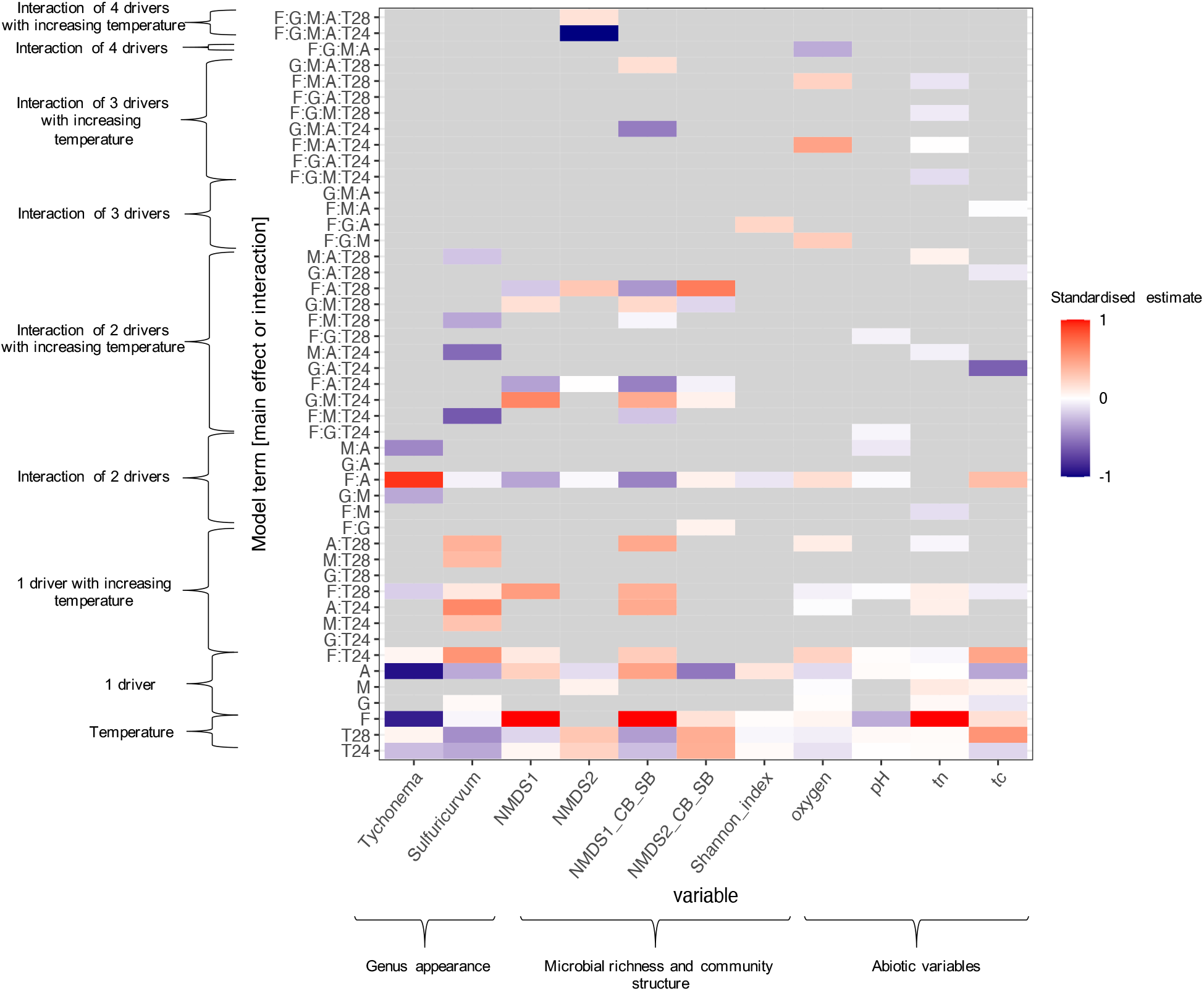
Summary of statistical analyses of all variables analysed. Eleven variables are shown in the columns of the heat map, (grouped by genus appearance, microbial richness and community structure and abiotic factors). Rows show the estimated coefficients of the single, one-way, two-way, three-way, and four-way interaction terms. Grey cells indicate a response variable and coefficient pairs for which the coefficients were not significantly different from zero (t-test p-value > 0.05), otherwise the diverging colour palette illustrates the direction of the influence by the driver or interaction of drivers (estimates of each variable were standardized by dividing by largest absolute value of the estimates in each variable). See the main text for the applied response variable transformations. tn = total nitrogen, tc= total carbon.

**Fig. 4.**
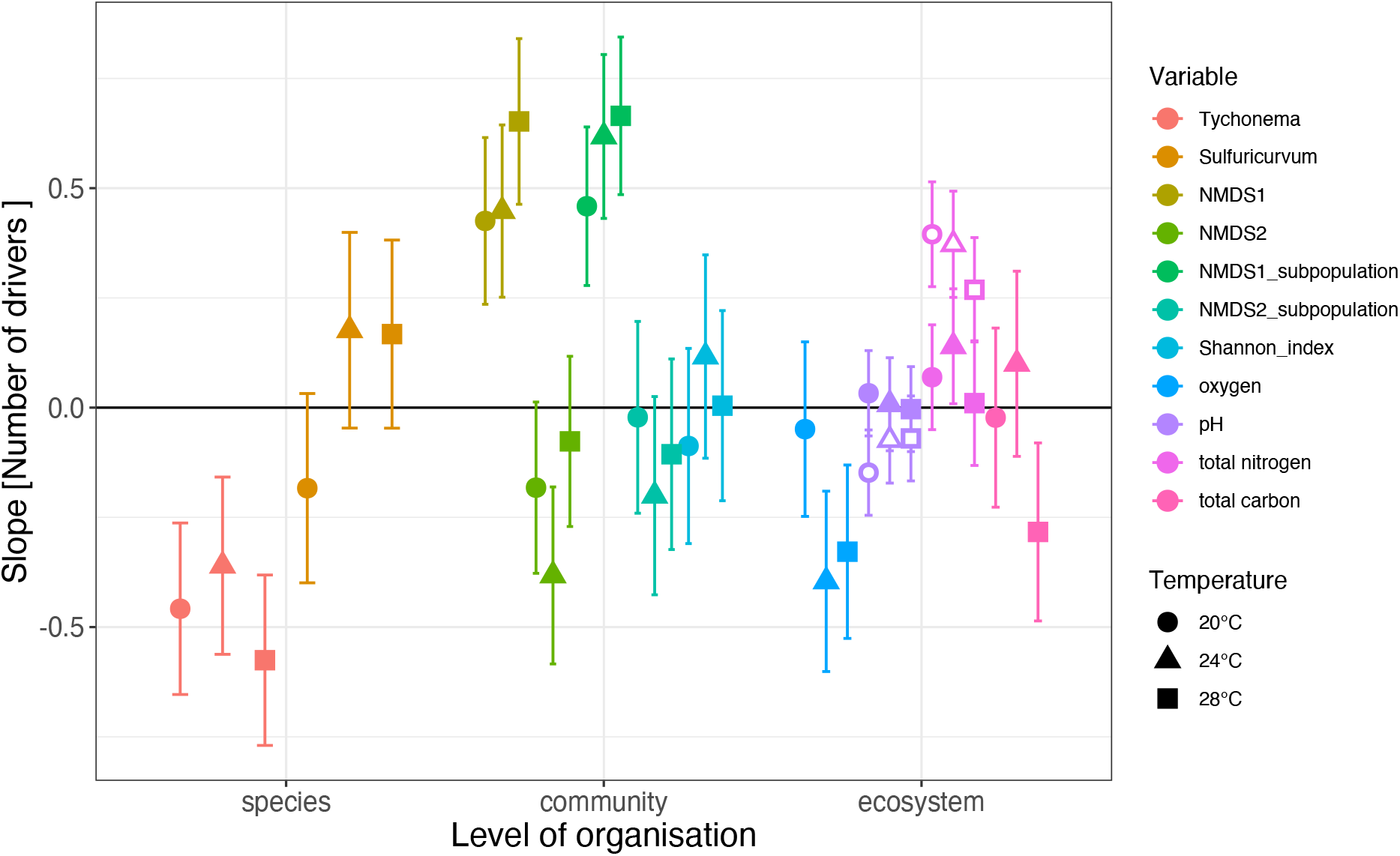
Effect size of number of drivers for each response variable at each temperature. Means and 95 % confidence intervals are shown. For the variables of pH and total nitrogen, fertilizer was used as a covariate due to its strong influence. Filled total nitrogen and pH labels: fertilizer was used as a driver. Empty total nitrogen and pH labels: fertilizer was excluded as driver. See the main text for the applied response variable transformations.

### Effects of combination and number of drivers on microbial community structure (based on NMDS)

Fertilizer addition was a key driver of microbial community composition (quantified via NMDS). (Fig. 2b+c, Supplementary Figure 5). Numerous driver combinations showed significant effects on NMDS score 1 and NMDS score 2 (Fig. 2c for NMDS1, Supplementary Fig. 4 for NMDS2, Fig. 3 for both), ranging from single to four-way effects. Different impact directions (positive or negative) can be identified depending on the driver combinations (Fig. 2c, Fig. 3). Please note that for clarity we also refer to NMDS scores with “negative” and “positive”, even though the direction is irrelevant for this measure. The three-way interaction of fertilizer+antibiotics+temperature on NMDS1 score is a good example how combination of drivers can change the impact direction (Fig. 5): When no fertilizer was present, the NMDS1 score was negative, while in presence of fertilizer, the NMDS1 score was positive. Additionally the combination of antibiotics and temperature changed the magnitude of the NMDS1 score. NMDS1 score was affected by number of drivers (F-statistic = 82.8, p-value < 0.001) (Fig. 2d, Fig. 4), and NMDS2 score was affected by both number of drivers (F-statistic = 12.7, p-value < 0.001) and temperature (F-statistic = 27.8, p-value < 0.001, Supplementary Figure 4b, Fig. 4). For both NMDS scores there was no interaction between number of drivers and temperature.

**Fig. 5.**
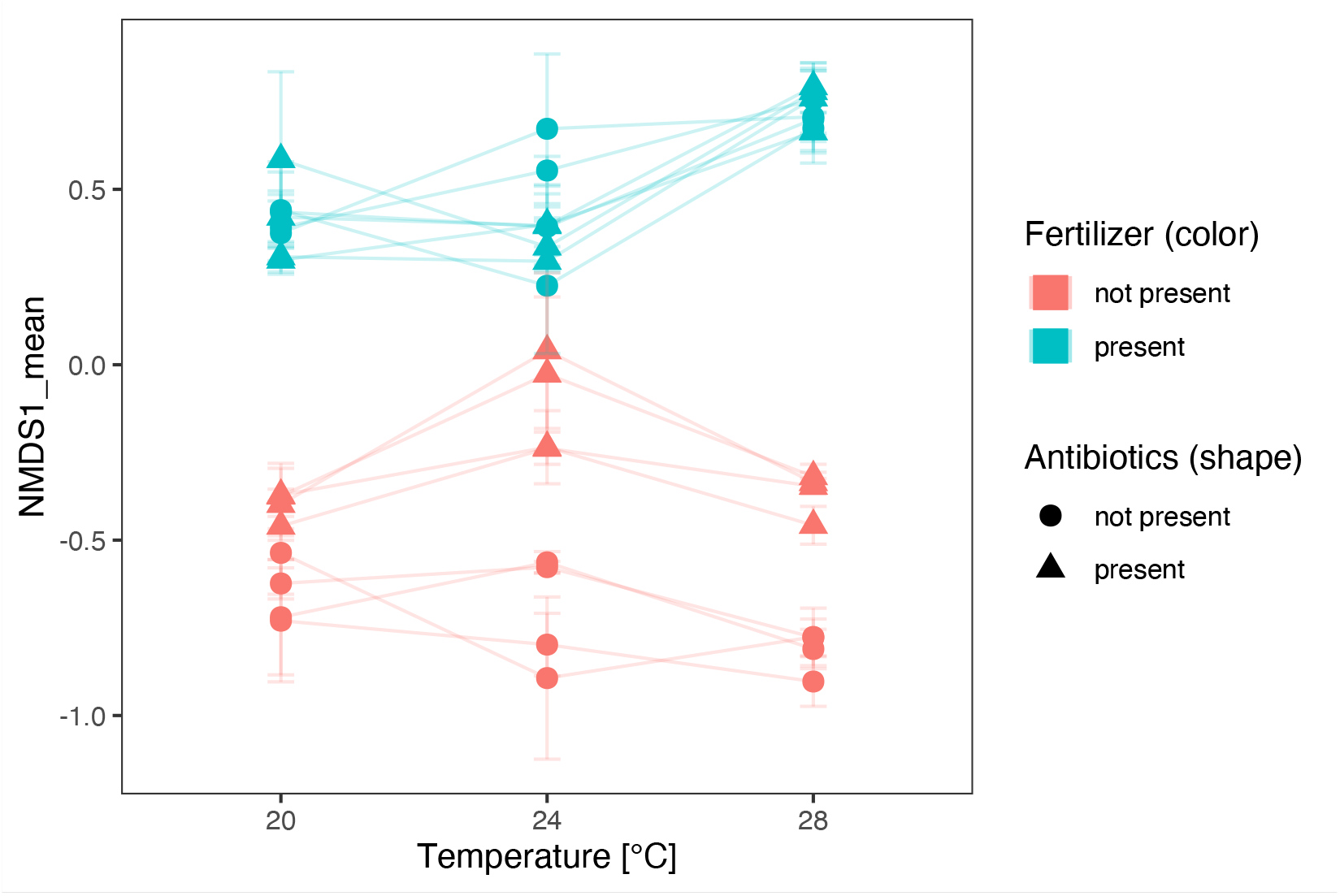
Visualization of the significant driver interactions of temperature, fertilizer and antibiotics on NMDS1. NMDS1 is affected by the presence of fertilizer, the presence of antibiotics and temperature, but also by the interactions of fertilizer+antibiotics and fertilizer+antibiotics+temperature. Means (n=5) are shown. Error bars represent standard error.

Analysis of the community of the subset of cyanobacteria, phototrophic sulfur bacteria and sulfate-reducing bacteria confirmed the trends observed for NMDS of all ASVs (Fig. 2e for NMDS1, Supplementary Fig. 4 for NMDS2, Fig. 3 for both), but showed slight differences on three-factorial and four-factorial driver combinations.

### Effects of combination and number of drivers on abiotic factors

Oxygen concentration was highly dependent on the driver combinations used, and various strong effects were identified (Fig. 2g), both positive and negative in direction. All drivers combined had a negative effect on oxygen concentration (F-statistic = 4.3, p-value = 0.040, non-standardised estimate -6.1 %). The number of drivers applied had a strong effect on the oxygen concentration at 24 °C and 28 °C (F-statistic = 18.8, p-value < 0.001), as well as temperature (F-statistic = 17.9, p-value < 0.001), and the interaction between temperature and number of drivers (F-statistic = 3.3, p-value = 0.040, Fig. 2h, Fig. 4).

The pH of the microcosms was influenced by numerous treatment combinations, but the strongest influences was identified as fertilizer addition, which decreased the pH from 8.5 to 6.5 (F-statistic = 3.2 × 103, p-value < 0.001, Fig. 2i). Driver combinations that included fertilizer addition had lower magnitude though were of the same sign (Fig. 3). Number of drivers had a negative effect on pH (F-statistic = 4.4, p-value = 0.037), with the clearest effect at 20 °C and no fertilizer used, in which case pH decreased the more drivers were applied (Fig. 2j, Fig. 4).

Effects on total nitrogen were driven by addition of fertilizer (F-statistic = 7.4 × 104, p-value < 0.001), but also numerous other combinations had a significant effect (Fig. 3, Supplementary Fig. 4). Furthermore, number of drivers strongly affected the total nitrogen concentration (F-statistic = 67.8, p-value < 0.001). Total carbon concentration was significantly affected by fertilizer (F-statistic = 31.9, p-value = 0.005), metal pollution (F-statistic = 9.9, p-value = 0.002) and temperature (F-statistic = 23.4, p-value < 0.001), as well as by antibiotics (F-statistic = 9.2, p-value = 0.003) and numerous combinations (Fig. 3, Supplementary Fig. 4). Furthermore, for the concentration of total carbon there was a significant interaction between number of drivers and temperature (F-statistic = 3.5, p-value = 0.032), with TC increasing with number of applied drivers at 28°C (Fig. 4, note that the response variable is the square inverse of total carbon).

## Discussion

Our study revealed compelling evidence of effects of the combination and the total number of drivers on taxa abundances, community composition, and ecosystem properties, and as such, that multiple drivers acting in combination can have important consequences across levels of ecological organisation (Rillig et al. 2019). Specifically, the addition of a driver, and/or temperature increase, could change the biological consequences of already applied drivers, even when the added driver applied alone had no effect (Fig. 3). We observed three trends here. The addition of a further driver could increase (e.g. F:A:T28), decrease (e.g. F:T28) or reverse the effect of the drivers which were already applied (e.g. F:A on *Tychonema* abundance, Fig. 6). This demonstrates clearly the need of analysing multiple drivers in all combinations. Furthermore, our study also confirmed that the number of drivers occurring together can influence microbial communities and thereby provide some hope that number of drivers can be used as a useful predictor, though individual driver effects can greatly contribute to the effect of number of drivers (Rillig et al. 2019). Furthermore, the effect of the number of drivers can be negative (e.g., *Tychonema* abundance) or positive (e.g. total nitrogen) and of different magnitudes, which highlights again the need of analysing various ecological response variables of species, microbial communities, and the whole ecosystem. While there may be some patterns across levels of organisation, e.g., the largest effects of number of drivers were observed at species and community levels of organisation (greater than 0.5 or less than -0.5 standardised effect size), the absence of strong patterns limits the scope for explanation and prediction of effect of multiple drivers across levels of organisation.

**Fig. 6.**
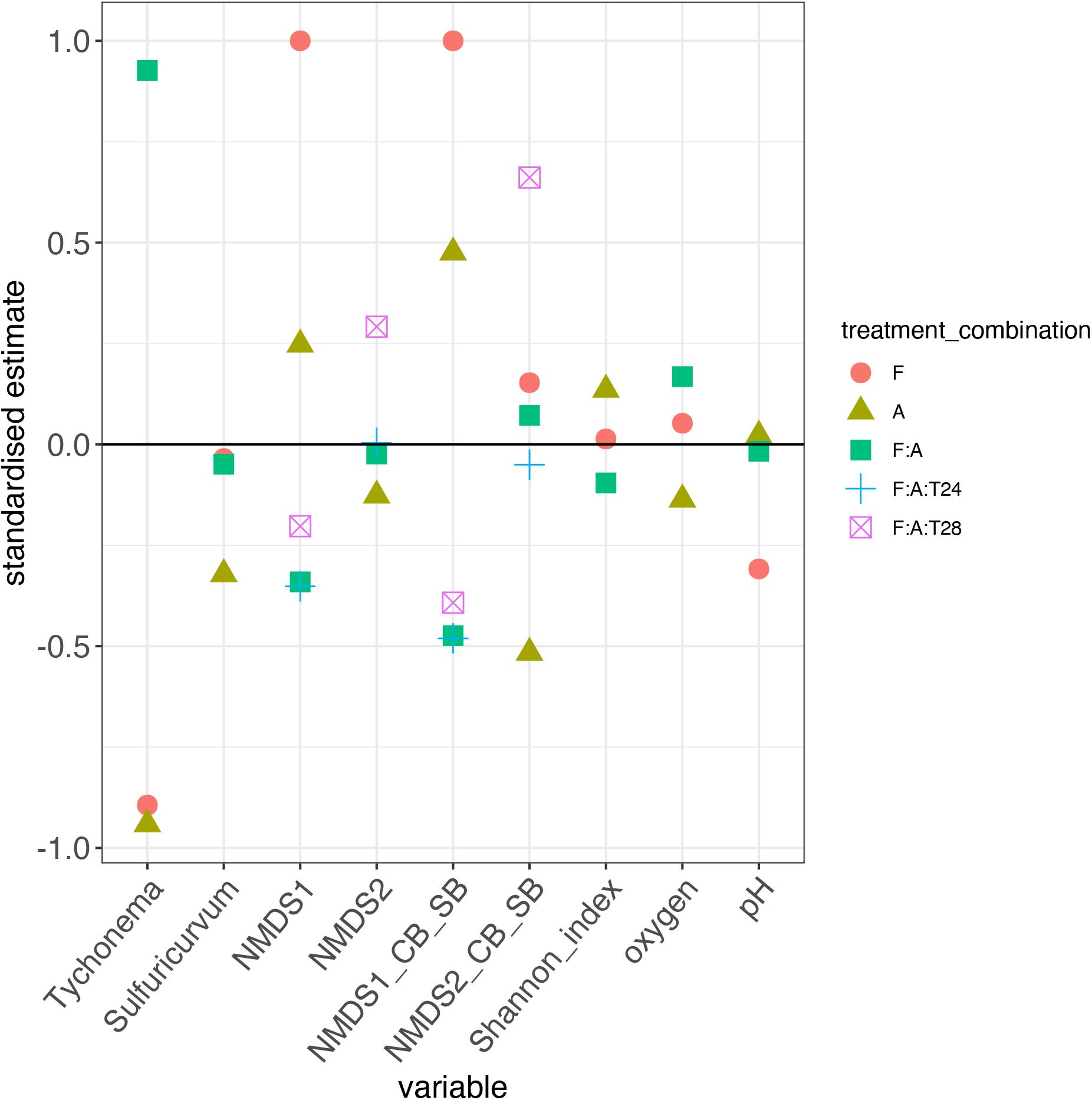
Effects of fertilizer, antibiotics and their combination at 20, 24, and 28 °C on ASV relative abundance, microbial composition, and ecosystem variables. Only significant treatments / treatment combinations are shown (F-test p value < 0.05). Total nitrogen and total carbon are not listed in this figure because they were not significantly affected by F, A, and F:A.

It is possible that effects of number of drivers is caused by multiple drivers acting in a similar fashion as a single driver of greater intensity. I.e. two drivers may be considered to typically have a higher effect because the overall intensity is higher. This would imply that drivers combine additively, or at least not completely substitutively. If this is the case, then some of the effects of number of drivers may not be so much number of drivers per se, but rather that total amount of driving being applied. Given that responses can be non-linear (and typically are), we can not exclude that some trends observed may simply be intensity effects.

A clear pattern in the results is of less apparent variability at 0 and at 4 drivers applied than with 1, 2, or 3 applied. This mainly because there are fewer combinations possible (1 combination possible to be precise: all or none), while at 1 to 3 drivers there are more combination of drivers possible, which leads to more variance. In contrast variance within driver combination is roughly invariant regardless of number of drivers. We are certain if we had a 5th stressor applied, we would see much more variation at 4 stressors applied, again because then we would have (5 choose 4)=5 possible stressor combinations.

While fertilizer, metal pollution and antibiotics had detectable effects when applied as single drivers, glyphosate did not. This finding is likely quite specific to the concentration of treatments used and should be regarded with caution. Nevertheless, it is even more interesting that when adding glyphosate to other drivers it did have an effect (see e.g., Shannon index, Fig. 3). Strong interactions were detected for antibiotics and fertilizer, and interestingly, these drivers revealed in combination different biological consequences compared to applied as a single driver, and temperature could also change the effects of the fertilizer-antibiotic driver duo again (Fig. 6).

The effect of temperature is in our view important to note, since it predicts that observed biological consequences caused by global change can change with on-going global warming. An increase of temperature can lead to new interactions among driver combinations (F:A with increasing temperature led again to significant affected variables) or can lead to significant effects that were not observed at 20 °C (G:M, G:M:24 and G:M:28 for NMDS1).

Our study also contributes to narrowing the gap of understanding of how drivers and their combination act across various levels of organisation. Our results revealed that individual taxa can be affected differently compared to abiotic factors of the ecosystem or the microbial community composition. Therefore, our study is in line with the findings of Galic et al. (2018) who reported differences of the influence of multiple drivers across levels of organisation. Interestingly, in our study, some treatment combinations, like fertilizer and antibiotics, showed significant influence across most variables and the three levels of organisation (except total nitrogen and total carbon), though even then the sign of the interaction term was sometimes negative and sometimes positive.

Effects of drivers, such as those we manipulated, are of course concentration-dependent. In this study, we used only two levels of driver concentration and added the driver in a single pulse treatment. More information about specific concentration thresholds and press disturbances are important to study and may clarify some observed trends, as well as the timing of the driver applied. In addition, responses of functional (metabolic) aspects of the aquatic microbial communities should be investigated in future studies, which could shed light into activated and deactivated pathways and enzymes used in microbial communities to react to changing environmental conditions.

One limitation of this study is that the background level of chemical stressor was not analysed prior to stressor addition. This information is important, since prior exposure to pollution would influent the responses to additional exposures. We expect that the small pond which was used to provide the water and sediment sample is low-nutrient loaded, and is due to its location largely free from herbicide, metal-, and antibiotic contamination. Nevertheless, this study showed clearly that combinations of stressors can lead to a new biological consequence, and this finding is very likely not impacted by any pre-exposure of drivers of the small pond.

One promising avenue for further research about species responses to multiple environmental drivers is to attempt to relate the patterns of response to features of the species, such as functional traits of the species. A relationship would reveal what traits of the species are determining how they respond to multiple drivers, and then give potential to predict those responses from species traits. This could be done for already present species, and also potentially invasive species. In studies such as ours, this would involve relating response patterns to characteristics of the microbial ASVs, and hence would require mapping of ASVs to trait information, which is not straightforward. Another option would be to use the 16S rRNA gene sequences to predict functional characteristics of the ASVs (e.g. (Ling et al. 2022).

In sum, our study confirmed the need to research the effects of multiple drivers on microbial communities and indicates further that a broad range of levels of organisation (species, community, ecosystem) should be analysed due to unique sets of effect and interactions across these and the different variables within them. Furthermore, our study showed that combination of specific drivers can change the biological consequence and direction compared to single drivers at all levels of organisation and that the effects of driver combinations can be modified by temperature. Overall, the lack of patterns in effect sizes across levels of organisation, and with respect to the number of drivers, represents lack of evidence for these to be useful in the prediction of the effects of combinations of global change drivers on ecological communities.

Finally, it is unclear how transferable our findings are to non-microbes or even to other types of microbial ecosystems. Microbes have some unique characteristics, such as mechanisms by which genetic material can move among individuals (horizontal gene transfer). There may be other important differences, such as strong interspecific interactions, particularly in closed experimental communities. Clearly what is required is a sufficient number of experiments, across a diverse enough collection of ecosystem and organism types, that involve at least four environmental drivers and observation of responses at population, community and ecosystem level, such that a formal meta-analysis is possible. This could be achieved a global network of such multi-stressor factorial experiments studying responses across levels of ecological organization.

## Acknowledgements

MS was funded by Forschungskredit of the UZH (FK-20-125). OLP was supported by the University of Zurich Research Priority Programmne in Global Change and Biodiversity. OLP and CZ were supported by Swiss National Science Foundation (Project 310030_188431). We thank the Predictive Ecology group for comments that greatly improved the project and this report of it. We thank the Functional Genomic Center Zurich for sequencing efforts.

## Author contribution

OLP and MS planned the experimental set-up. MS performed all experiments in the lab. MS performed the up- and downstream bioinformatics. MS, OLP and UD performed the data analysis. UD supported the data transformation and statistical models. YC provided technical support and performed the TOC analysis. XZ performed the DNA extractions and provided technical support. OLP, MS and UD drafted the manuscript. All authors confirmed the final version of the manuscript.

## Supplementary Information

**Supplementary Fig. 1.**
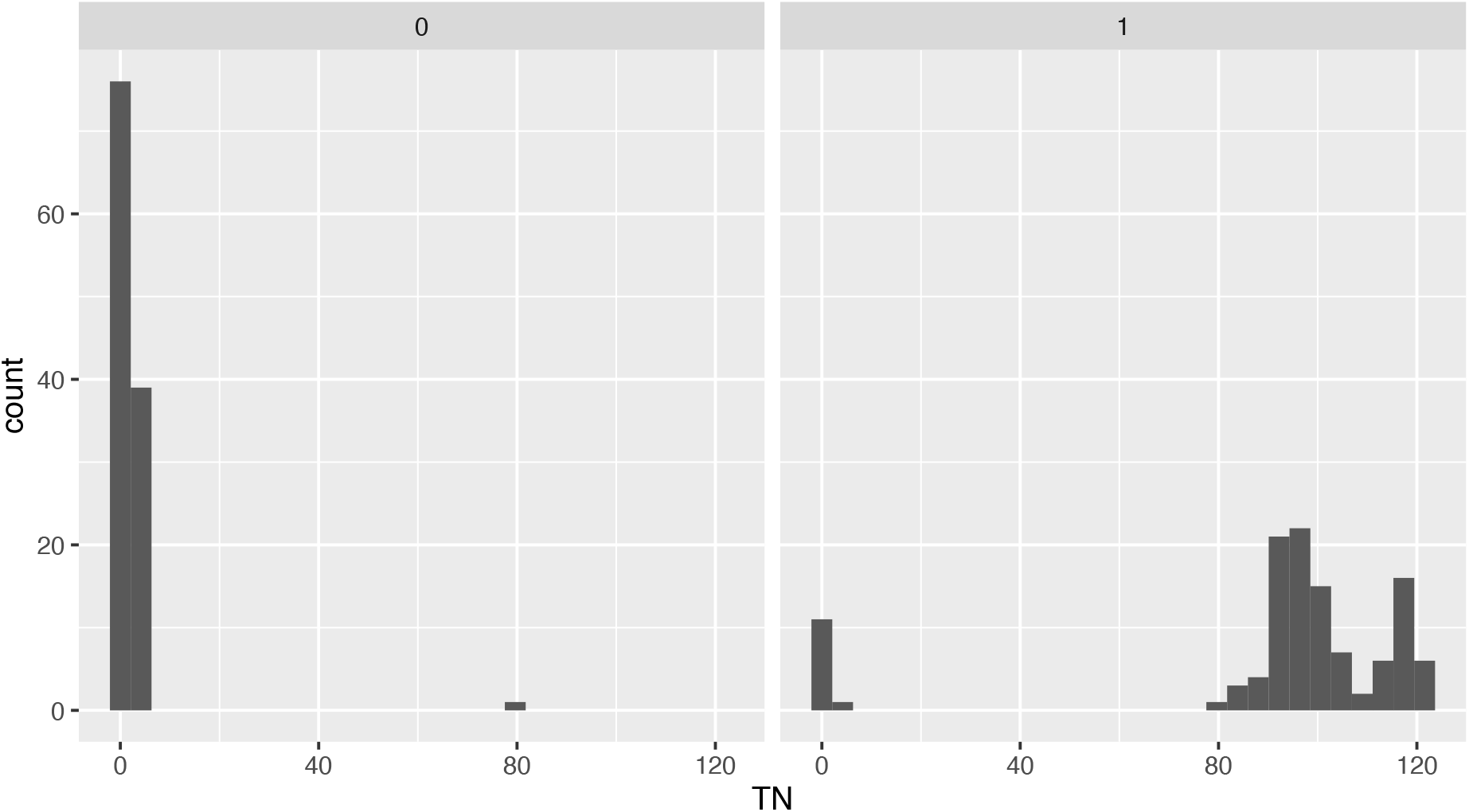
Exclusion of data of the total nitrogen measurement. There was one sample in which high amounts of nitrogen were detected when no fertilizer had been added and 12 samples in which no nitrogen was detected despite the fact that fertilizer had been added to them. Both of these two cases are not possible and were due to a faulty behavior of the measurement apparatus. Based on the measured nitrogen values in the other samples and their distribution we are confident that the measurement problems did not extend to them and there is thus no reason to doubt their validity.

**Supplementary Fig. 2.**
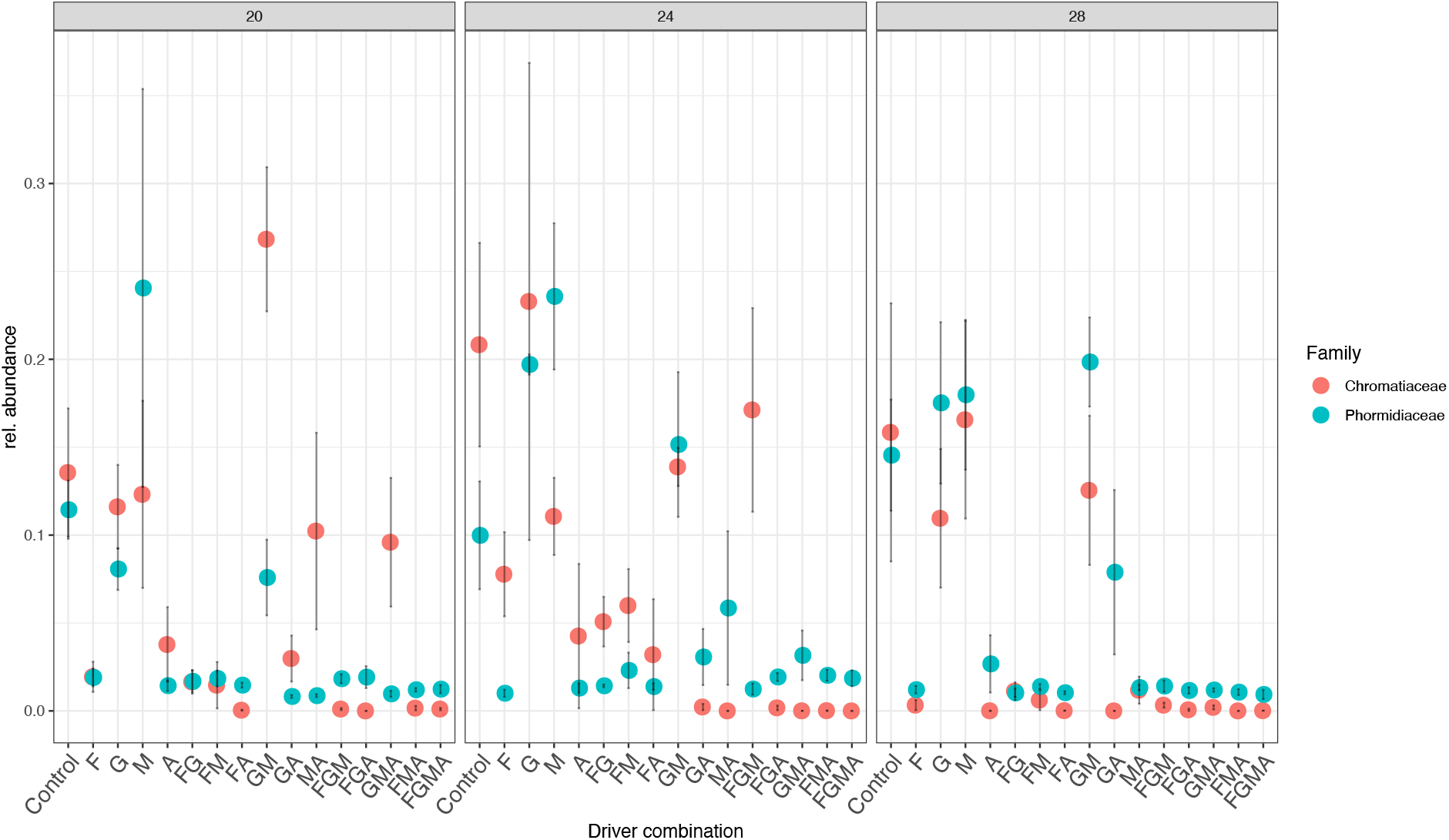
Relative abundance of *Chromatiaceae* and *Phormidiaceae* in the microcosm depending on driver combination and temperature. Mean is shown by the circle, standard errors are shown, n=5)

**Supplementary Fig. 3.**
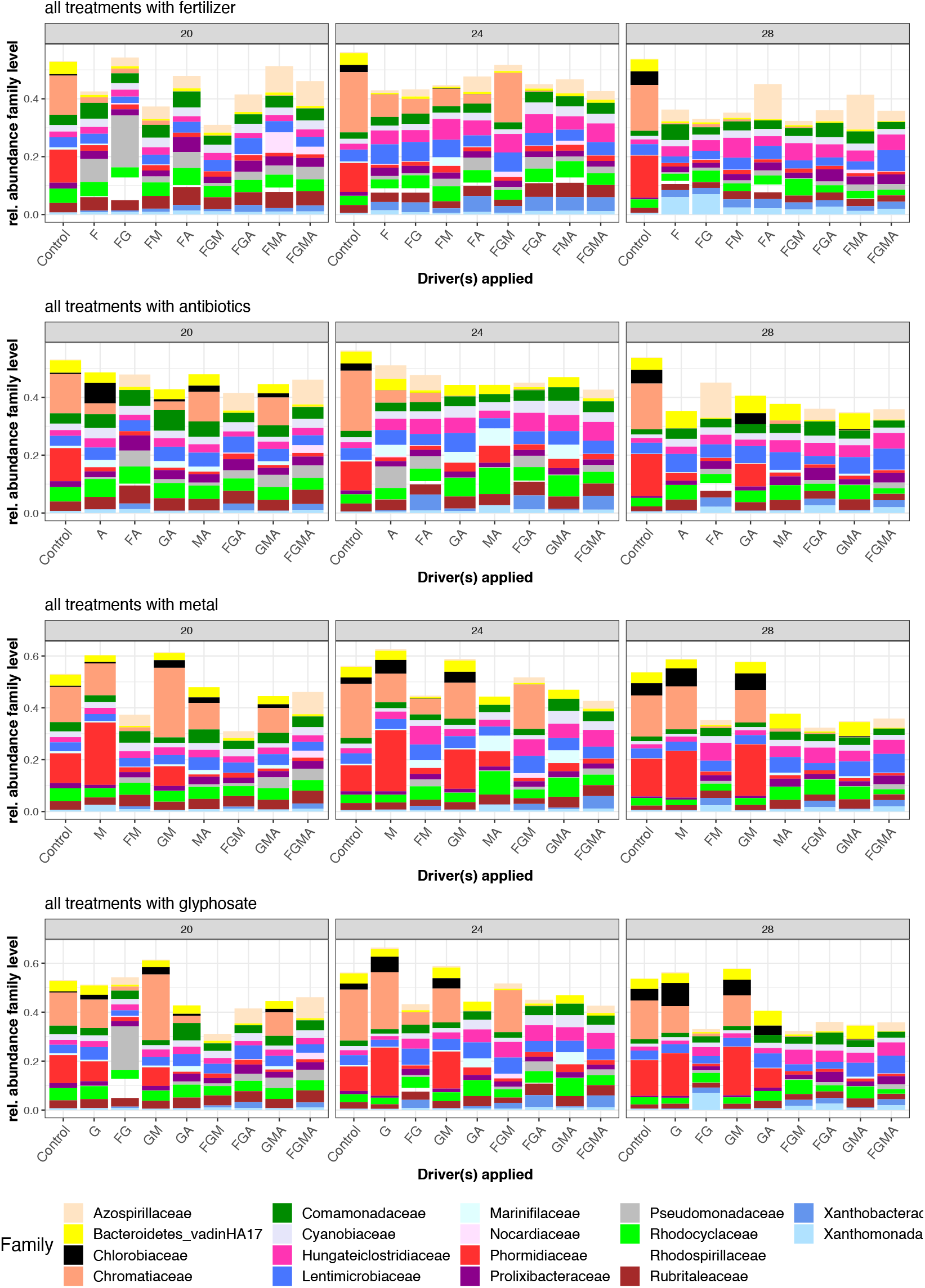
Effects of drivers and driver combinations on microbial community composition on family level (mean per 5 replicates) per driver category. Taxa with rel. abundance > 5 % shown.

**Supplementary Fig. 4.**
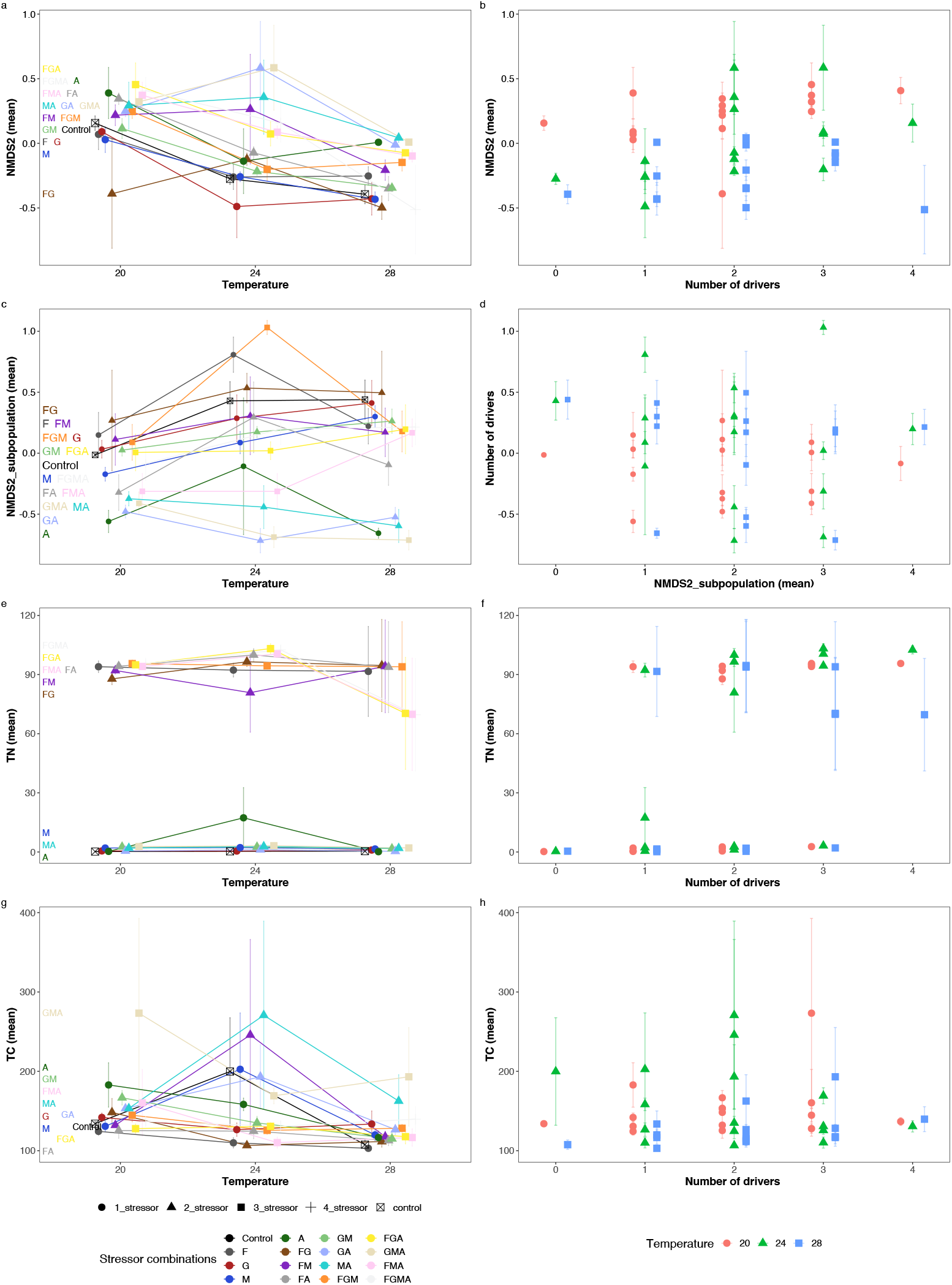
Effects of driver combination and number of drivers on temperature applied for microbial community and ecosystem variables. NMDS2 scores varying with (a) temperature and driver combination and (b) temperature and number of drivers. NMDS2_subpopulation scores varying with (c) temperature and driver combination and (d) temperature and number of drivers. Total nitrogen concentration varying with (e) temperature and driver combination and (f) temperature and number of drivers. Total carbon concentration varying with (g) temperature and driver combination and (h) temperature and number of drivers. Standard errors are shown (n = 5 replicates).

**Supplementary Fig. 5.**
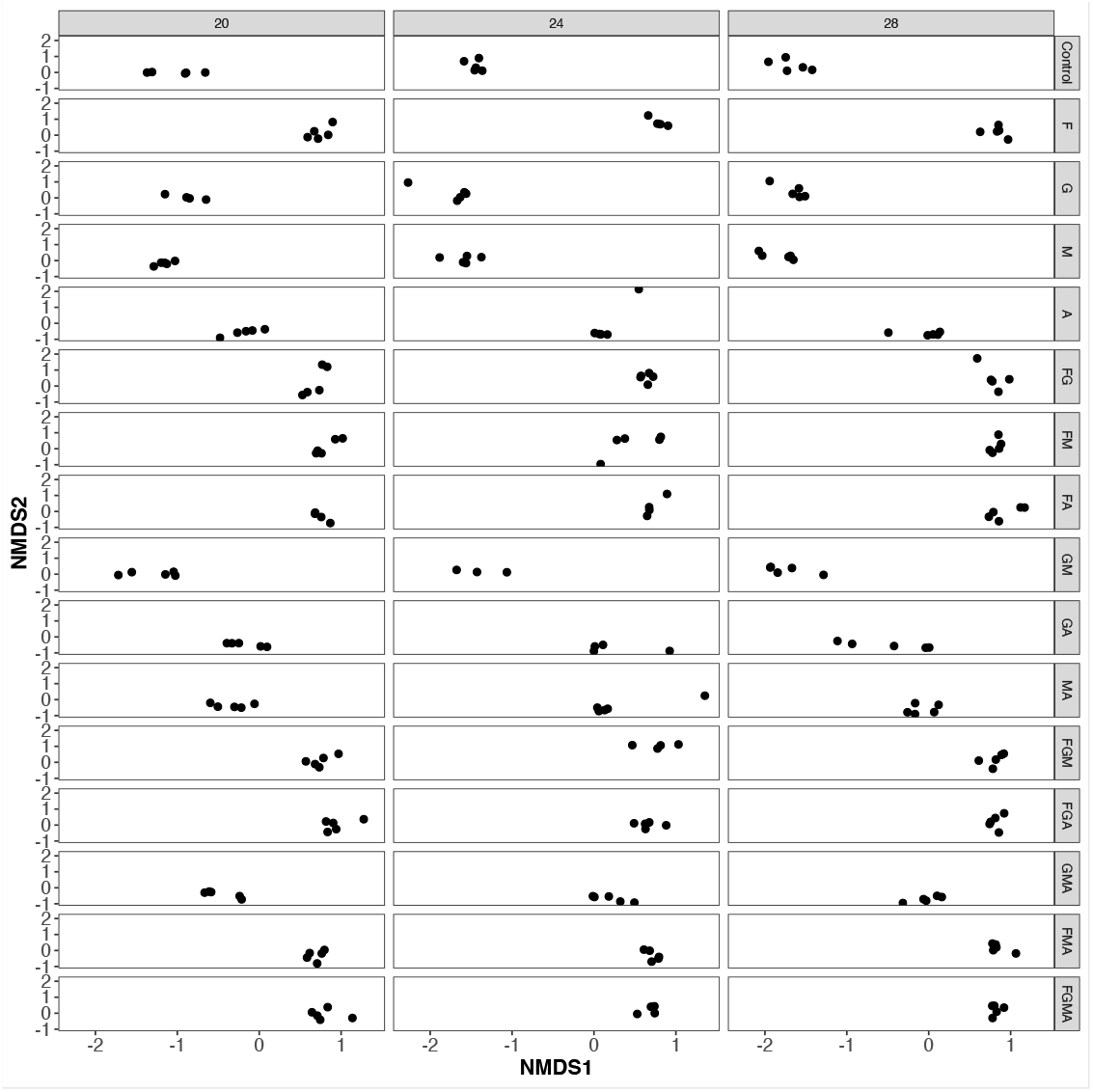
Beta-diversity based on NMDS analysis (Bray-Curtis) in relation to temperature and driver combinations. Stress is 0.16

**Supplementary Table 1.** Number of reads in each replicate.

| Combinations | total_reads | Replicate | Temperature |
| --- | --- | --- | --- |
| Control | 7364 | Replicate 1 | 24 |
| Control | 7046 | Replicate 1 | 28 |
| Control | 4659 | Replicate 1 | 20 |
| F | 9933 | Replicate 1 | 28 |
| F | 5912 | Replicate 1 | 20 |
| G | 265 | Replicate 1 | 24 |
| G | 8662 | Replicate 1 | 28 |
| M | 2695 | Replicate 1 | 24 |
| M | 8202 | Replicate 1 | 28 |
| M | 263 | Replicate 1 | 20 |
| A | 4195 | Replicate 1 | 24 |
| A | 7550 | Replicate 1 | 28 |
| A | 5276 | Replicate 1 | 20 |
| FG | 10815 | Replicate 1 | 24 |
| FG | 8945 | Replicate 1 | 28 |
| FM | 5660 | Replicate 1 | 20 |
| FM | 284 | Replicate 1 | 24 |
| FM | 8857 | Replicate 1 | 28 |
| FG | 4639 | Replicate 1 | 20 |
| FA | 4449 | Replicate 1 | 24 |
| FA | 8835 | Replicate 1 | 28 |
| FA | 3816 | Replicate 1 | 20 |
| GM | 7678 | Replicate 1 | 28 |
| GM | 6273 | Replicate 1 | 20 |
| GA | 7469 | Replicate 1 | 28 |
| GA | 6229 | Replicate 1 | 20 |
| MA | 10849 | Replicate 1 | 24 |
| MA | 11268 | Replicate 1 | 28 |
| MA | 18182 | Replicate 1 | 20 |
| FGM | 996 | Replicate 1 | 24 |
| FGM | 6189 | Replicate 1 | 28 |
| FGM | 6489 | Replicate 1 | 20 |
| FGA | 2766 | Replicate 1 | 24 |
| FGA | 8313 | Replicate 1 | 28 |
| FGA | 6763 | Replicate 1 | 20 |
| GMA | 9612 | Replicate 1 | 24 |
| GMA | 8460 | Replicate 1 | 28 |
| GMA | 7501 | Replicate 1 | 20 |
| FMA | 11304 | Replicate 1 | 24 |
| FMA | 8822 | Replicate 1 | 28 |
| FMA | 6340 | Replicate 1 | 20 |
| FGMA | 10657 | Replicate 1 | 24 |
| FGMA | 14034 | Replicate 1 | 28 |
| FGMA | 5819 | Replicate 1 | 20 |
| Control | 9426 | Replicate 2 | 24 |
| Control | 7438 | Replicate 2 | 28 |
| Control | 2301 | Replicate 2 | 20 |
| F | 5416 | Replicate 2 | 24 |
| F | 9076 | Replicate 2 | 28 |
| F | 6040 | Replicate 2 | 20 |
| G | 7175 | Replicate 2 | 24 |
| G | 8350 | Replicate 2 | 28 |
| G | 5573 | Replicate 2 | 20 |
| M | 7295 | Replicate 2 | 24 |
| M | 6408 | Replicate 2 | 28 |
| M | 7580 | Replicate 2 | 20 |
| A | 7606 | Replicate 2 | 24 |
| A | 7916 | Replicate 2 | 28 |
| A | 13579 | Replicate 2 | 20 |
| FG | 2733 | Replicate 2 | 24 |
| FG | 9212 | Replicate 2 | 28 |
| FG | 5454 | Replicate 2 | 20 |
| FM | 7872 | Replicate 2 | 24 |
| FM | 6398 | Replicate 2 | 28 |
| FM | 4005 | Replicate 2 | 20 |
| FA | 10016 | Replicate 2 | 24 |
| FA | 7423 | Replicate 2 | 28 |
| FA | 5710 | Replicate 2 | 20 |
| GM | 11100 | Replicate 2 | 24 |
| GM | 8294 | Replicate 2 | 28 |
| GM | 7579 | Replicate 2 | 20 |
| GA | 10206 | Replicate 2 | 24 |
| GA | 7078 | Replicate 2 | 28 |
| GA | 8674 | Replicate 2 | 20 |
| MA | 13163 | Replicate 2 | 24 |
| MA | 7927 | Replicate 2 | 28 |
| MA | 7229 | Replicate 2 | 20 |
| FGM | 10680 | Replicate 2 | 24 |
| FGM | 8434 | Replicate 2 | 28 |
| FGM | 8146 | Replicate 2 | 20 |
| FGA | 12236 | Replicate 2 | 24 |
| FGA | 8018 | Replicate 2 | 28 |
| FGA | 7086 | Replicate 2 | 20 |
| GMA | 10598 | Replicate 2 | 24 |
| GMA | 8885 | Replicate 2 | 28 |
| GMA | 6503 | Replicate 2 | 20 |
| FMA | 5447 | Replicate 2 | 24 |
| FMA | 9021 | Replicate 2 | 28 |
| FMA | 12484 | Replicate 2 | 20 |
| FGMA | 2177 | Replicate 2 | 24 |
| FGMA | 7974 | Replicate 2 | 28 |
| FGMA | 4190 | Replicate 2 | 20 |
| Control | 10159 | Replicate 3 | 24 |
| Control | 8105 | Replicate 3 | 28 |
| Control | 3981 | Replicate 3 | 20 |
| F | 9959 | Replicate 3 | 24 |
| F | 7459 | Replicate 3 | 28 |
| F | 6840 | Replicate 3 | 20 |
| G | 10556 | Replicate 3 | 24 |
| G | 8158 | Replicate 3 | 28 |
| G | 6546 | Replicate 3 | 20 |
| M | 11471 | Replicate 3 | 24 |
| M | 12913 | Replicate 3 | 28 |
| M | 8389 | Replicate 3 | 20 |
| A | 11192 | Replicate 3 | 24 |
| A | 10260 | Replicate 3 | 28 |
| A | 6407 | Replicate 3 | 20 |
| FG | 12182 | Replicate 3 | 24 |
| FG | 10109 | Replicate 3 | 28 |
| FG | 7141 | Replicate 3 | 20 |
| FM | 9788 | Replicate 3 | 24 |
| FM | 8356 | Replicate 3 | 28 |
| FM | 8224 | Replicate 3 | 20 |
| FA | 11486 | Replicate 3 | 24 |
| FA | 8163 | Replicate 3 | 28 |
| FA | 5961 | Replicate 3 | 20 |
| GM | 9096 | Replicate 3 | 24 |
| GM | 8953 | Replicate 3 | 28 |
| GM | 10665 | Replicate 3 | 20 |
| GA | 2887 | Replicate 3 | 24 |
| GA | 7543 | Replicate 3 | 28 |
| GA | 5011 | Replicate 3 | 20 |
| MA | 21214 | Replicate 3 | 24 |
| MA | 7612 | Replicate 3 | 28 |
| MA | 6836 | Replicate 3 | 20 |
| FGM | 10304 | Replicate 3 | 24 |
| FGM | 8851 | Replicate 3 | 28 |
| FGM | 7718 | Replicate 3 | 20 |
| FGA | 11639 | Replicate 3 | 24 |
| FGA | 10839 | Replicate 3 | 28 |
| FGA | 7296 | Replicate 3 | 20 |
| GMA | 14752 | Replicate 3 | 24 |
| GMA | 10593 | Replicate 3 | 28 |
| GMA | 8784 | Replicate 3 | 20 |
| FMA | 12812 | Replicate 3 | 24 |
| FMA | 11821 | Replicate 3 | 28 |
| FMA | 9476 | Replicate 3 | 20 |
| FGMA | 12513 | Replicate 3 | 24 |
| FGMA | 12886 | Replicate 3 | 28 |
| FGMA | 6091 | Replicate 3 | 20 |
| Control | 11749 | Replicate 4 | 24 |
| Control | 9920 | Replicate 4 | 28 |
| Control | 5989 | Replicate 4 | 20 |
| F | 10527 | Replicate 4 | 24 |
| F | 9335 | Replicate 4 | 28 |
| F | 7371 | Replicate 4 | 20 |
| G | 7547 | Replicate 4 | 24 |
| G | 8513 | Replicate 4 | 28 |
| G | 10282 | Replicate 4 | 20 |
| M | 3827 | Replicate 4 | 24 |
| M | 6486 | Replicate 4 | 28 |
| M | 5361 | Replicate 4 | 20 |
| A | 9307 | Replicate 4 | 24 |
| A | 8114 | Replicate 4 | 28 |
| A | 5193 | Replicate 4 | 20 |
| FG | 11085 | Replicate 4 | 24 |
| FG | 11630 | Replicate 4 | 28 |
| FG | 6776 | Replicate 4 | 20 |
| FM | 10109 | Replicate 4 | 24 |
| FM | 8542 | Replicate 4 | 28 |
| FM | 7170 | Replicate 4 | 20 |
| FA | 11529 | Replicate 4 | 24 |
| FA | 9826 | Replicate 4 | 28 |
| FA | 5165 | Replicate 4 | 20 |
| GM | 13030 | Replicate 4 | 24 |
| GM | 10250 | Replicate 4 | 28 |
| GM | 8414 | Replicate 4 | 20 |
| GA | 12392 | Replicate 4 | 24 |
| GA | 9148 | Replicate 4 | 28 |
| GA | 7635 | Replicate 4 | 20 |
| MA | 12252 | Replicate 4 | 24 |
| MA | 8859 | Replicate 4 | 28 |
| MA | 7882 | Replicate 4 | 20 |
| FGM | 11554 | Replicate 4 | 24 |
| FGM | 10022 | Replicate 4 | 28 |
| FGM | 5482 | Replicate 4 | 20 |
| ori | 7942 | ori_2 | 19 |
| FGA | 7658 | Replicate 4 | 24 |
| FGA | 9903 | Replicate 4 | 28 |
| FGA | 4233 | Replicate 4 | 20 |
| GMA | 1965 | Replicate 4 | 24 |
| GMA | 7948 | Replicate 4 | 28 |
| GMA | 4641 | Replicate 4 | 20 |
| FMA | 5334 | Replicate 4 | 24 |
| FMA | 7455 | Replicate 4 | 28 |
| FMA | 3332 | Replicate 4 | 20 |
| FGMA | 11071 | Replicate 4 | 24 |
| FGMA | 8356 | Replicate 4 | 28 |
| FGMA | 5910 | Replicate 4 | 20 |
| Control | 13077 | Replicate 5 | 24 |
| Control | 7919 | Replicate 5 | 28 |
| Control | 6401 | Replicate 5 | 20 |
| F | 8301 | Replicate 5 | 24 |
| F | 7802 | Replicate 5 | 28 |
| F | 5801 | Replicate 5 | 20 |
| G | 11899 | Replicate 5 | 24 |
| G | 8686 | Replicate 5 | 28 |
| G | 2745 | Replicate 5 | 20 |
| M | 11401 | Replicate 5 | 24 |
| M | 8270 | Replicate 5 | 28 |
| M | 2541 | Replicate 5 | 20 |
| A | 7766 | Replicate 5 | 24 |
| A | 7256 | Replicate 5 | 28 |
| A | 497 | Replicate 5 | 20 |
| FG | 12676 | Replicate 5 | 24 |
| FG | 8439 | Replicate 5 | 28 |
| FG | 15008 | Replicate 5 | 20 |
| ori | 13295 | ori_5 | 19 |
| ori | 3919 | ori_3 | 19 |
| FM | 8865 | Replicate 5 | 24 |
| FM | 11599 | Replicate 5 | 28 |
| FM | 4819 | Replicate 5 | 20 |
| FA | 8044 | Replicate 5 | 28 |
| GM | 7221 | Replicate 5 | 28 |
| GM | 504 | Replicate 5 | 20 |
| GA | 229 | Replicate 5 | 24 |
| GA | 7704 | Replicate 5 | 28 |
| GA | 8461 | Replicate 5 | 20 |
| MA | 30 | Replicate 5 | 24 |
| MA | 8202 | Replicate 5 | 28 |
| MA | 2288 | Replicate 5 | 20 |
| FGM | 9054 | Replicate 5 | 28 |
| FGM | 3240 | Replicate 5 | 20 |
| FGA | 8673 | Replicate 5 | 24 |
| FGA | 9333 | Replicate 5 | 28 |
| FGA | 392 | Replicate 5 | 20 |
| GMA | 488 | Replicate 5 | 24 |
| GMA | 8907 | Replicate 5 | 28 |
| GMA | 6931 | Replicate 5 | 20 |
| FMA | 8137 | Replicate 5 | 24 |
| FMA | 9209 | Replicate 5 | 28 |
| FMA | 4630 | Replicate 5 | 20 |
| FGMA | 8672 | Replicate 5 | 28 |
| FGMA | 4883 | Replicate 5 | 20 |
| ori | 10488 | ori_6 | 19 |

